# Quality control of variant peptides identified through proteogenomics- catching the (un)usual suspects

**DOI:** 10.1101/2023.05.31.542998

**Authors:** Anurag Raj, Suruchi Aggarwal, Amit Kumar Yadav, Debasis Dash

**Affiliations:** G. N. Ramachandran Knowledge Centre for Genomics Informatics, CSIR – Institute of Genomics and Integrative Biology, New Delhi, India; Academy of Scientific and Innovative Research (AcSIR), Ghaziabad– 201002, India; Computational and Mathematical Biology Centre (CMBC), Translational Health Science and Technology Institute, NCR Biotech Science Cluster, 3rd Milestone, Faridabad, Haryana, India

**Keywords:** Proteogenomics, PgxSAVy, Variant peptides, Proteoforms, SAAVs, SAPs, Mass spectrometry, Proteomics, mutations, SNPs, False discovery rate, Quality assessment

## Abstract

Variant peptides resulting from translation of single nucleotide polymorphisms (SNPs) can lead to aberrant or altered protein functions and thus hold translational potential for disease diagnosis, therapeutics and personalized medicine. Variant peptides detected by proteogenomics are fraught with high number of false positives. Class-specific FDR along with ad-hoc post-search filters have been employed to tackle this issue, but there is no uniform and comprehensive approach to assess variant quality. These protocols are mostly manual or tedious, and not accessible across labs. We present a software tool, PgxSAVy, for the quality control of variant peptides. PgxSAVy provides a rigorous framework for quality control and annotations of variant peptides on the basis of (i) variant quality, (ii) isobaric masses, and (iii) disease annotation. PgxSAVy was able to segregate true and false variants with 98.43% accuracy on simulated data. We then used ∼2.8 million spectra (PXD004010 and PXD001468) and identified 12,705 variant PSMs, of which PgxSAVy evaluated 3028 (23.8%), 1409 (11.1%) and 8268 (65.1%) as confident, semi-confident and doubtful respectively. PgxSAVy also annotates the variants based on their pathogenicity and provides support for assisted manual validation. In these datasets, it identified previously found variants as well some novel variants not seen in original studies. The confident variants identified the importance of mutations in glycolysis and gluconeogenesis pathways in Alzheimer’s disease. The analysis of proteins carrying variants can provide fine granularity in discovering important pathways. PgxSAVy will advance personalized medicine by providing a comprehensive framework for quality control and prioritization of proteogenomics variants.

**Availability:** PgxSAVy is freely available at https://github.com/anuragraj/PgxSAVy

**Key Points:**

- Variant peptide in proteogenomics have high rates of false positives
- class-specific FDR is not sufficiently effective, and tedious manual filtering is not scalable
- We developed PgxSAVy for automated quality control and disease annotation of variant peptides from proteogenomics search results
- PgxSAVy was validated using simulation data and manually annotated variant PSMs
- Independent application on large datasets on Alzheimer’s and HEK cell lines demonstrated that PgxSAVy discovered known and novel mutations with important biological roles.

**Graphical Abstract:** 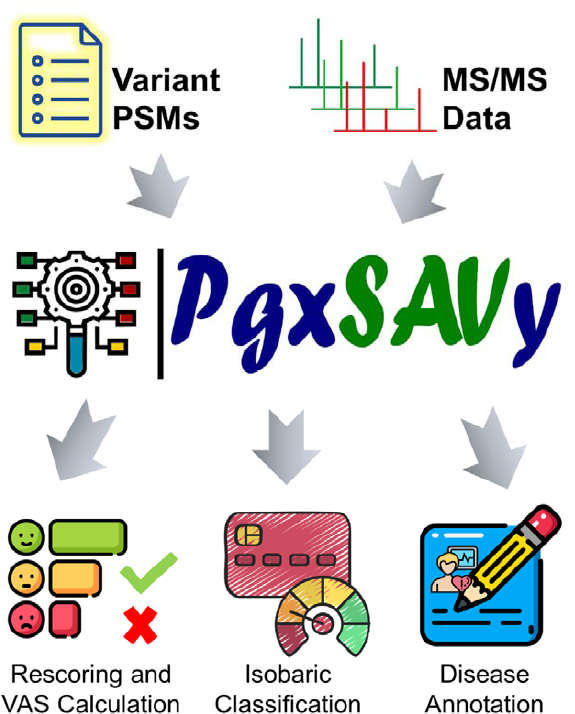

## Introduction

Single nucleotide polymorphisms (SNPs) play an important role in defining the health and disease status of an organism. Single amino acid variations (SAVs) caused due to SNPs can alter the structure, interactions or activity of the corresponding proteins [1,2]. Proteogenomics is a powerful approach to mine mass spectrometry data for identification and characterization of variants. Many variants of biological importance have been discovered in diseases like cancers, Alzheimer’s, Parkinson’s, diabetes and cardiovascular diseases [3–8]. Integrated analysis of multi-omics data can immensely benefit personalized medicine by proteogenomics analysis [9,10], which can directly connect patient-specific genomic variants and their protein level translation products. Thus, proteogenomics can provide important information for personalized medicine, by connecting disease susceptibility, diagnosis and response to drugs based on their associated variants [11,12].

In shotgun proteomics, the experimental MS/MS spectra are generated from fragmentation of tryptic peptides and matched against theoretical MS/MS spectra obtained from a reference protein database digested *in silico*. The database search tools such as X!Tandem [13], OMSSA [14], MSGF+ [15] etc. can identify peptides and their known modifications. The single amino acid variants (SAVs) are usually not present in the database and therefore not identified. The proteogenomics search against 3-frame translated transcriptome or 6-frame translated genome are best suited for variant peptide detection [16–20], but often contain many false positives. The statistical validation is performed by a target-decoy (TD) based false discovery rate (FDR) control, in which the data is searched simultaneously in a reference (target) database and its reversed-sequence (decoy) database. The FDR is estimated by ratio of decoy/target at any given threshold to control error rates [21–23].

Since the proteogenomics databases tend to be humongous in size, there is a higher-than-usual chances of false hits. To identify variants, usually a custom variant database is used which is much larger in size than a proteogenomics database [24]. Thus, it is also fraught with high rates of error that the target-decoy (TD) based FDR often fails to control [23,25]. This reduced sensitivity and increased rate of false positives pose major challenges in variant identification and are highlighted in recent articles [25–29]. Most proteogenomic studies calculate a global false discovery rate for all peptide identifications, including known, novel and variant peptides [30–32]. Some studies have recommended a higher threshold of FDR for novel peptides and implemented stringent filters or class-specific FDR (cFDR) [33]. Even with stricter thresholds, the problem remains recalcitrant as demonstrated in several studies [34–36]. Therefore, a comprehensive identification of the sequence variants of the proteins is still a formidable challenge [35].

The process of confidence assessment requires additional evaluation and filtering criteria after the database search and FDR estimation. Li et al. developed a modified FDR estimation workflow for evaluation of variant peptides, but also acknowledged that many false variants still remained in the data [37]. It is also required that the variant peptides should score higher than their wild type sequences. Another study proposed SpectrumAI tool to verify SAV peptides by requiring variant-event ions to directly support the residue substitution in MS/MS spectra [38]. However, it only checks variant amino acid peak but does not use intensity information, neither does it compare against the wildtype peptides. A comprehensive set of recommendations was proposed for variant peptide search and evaluation, in which the authors employed multi-algorithm searches and a split target-decoy approach (same as class-specific FDR) coupled with various post-search filters to remove false identifications [36]. This included isobaric checks to test if variant can be explained by other isobaric amino acids, PTM mass shift, reference proteome mapping to detect indistinguishable peptides and protein abundance. Similarly, the post-examination of variant peptides by applying filters and checking criteria like isobaric substitutions has been proposed recently [35] with the demonstration that standard FDR estimation was not sufficient in controlling variant false positives.

However, these assessment criteria and recommendations, although useful, are not amenable for automated verification of variant peptides. Also, since these recommendations are not adopted in a uniform manner across different labs, the implementations can differ widely. Due to lack of a standard tool, it either requires advanced bioinformatics skills or manual verification, neither of which is complete or comprehensive. Variant quality evaluation, thus requires robust and objective scoring system implemented as an automated pipeline. SAVControl [34] is a promising tool that filters variant peptides based on subgroup FDR. It additionally checks for variant amino acid position in the peptide by mapping given mass shift and residue against UniMod counterparts. It subsequently provides a classification based on mapping of site determined by PTMiner [39] with UniMod but does not perform in-depth check of variant quality.

Using stage-specific searches for the wild type peptides followed by variant peptides for unassigned spectra, helps in reducing false variants [40]. Some studies have also used multiple search engines to improve PSM identification rate by harnessing complementarity between search engines [36]. Measures like spectral match quality, intensity coverage and b/y ion counts are often employed to weed out false variants in manual verification. An easy to calculate variant quality measure that is implemented in a search-engine-agnostic pipelines is an important requirement or design goal to facilitate quality control in variant proteogenomics.

With these design goals and challenges in mind, we developed PgxSAVy tool that first rescores the variant peptides using a modified version of MassWiz score [41], and calculates *variant ambiguity score* (VAS) as a universally applicable variant scoring method. Here, we posited that variant peptide rescoring and integration of various features for PSM quality, variant event and global search results (i.e., PSMs count per peptides and/or search engine count) together into VAS can reduce false variants. These features add value in assessing the quality of variant peptide above and beyond TD based FDR, multiple search engines and cFDR. We have integrated the tedious step of manual validation, isobaric checks and available pathogenic/disease related information for human variant peptides into a seamless, standardized and easy-to-use tool. We have demonstrated the accuracy and utility using simulated spectra as well as large-scale data from Alzheimer’s disease and HEK293 cell lines. This will advance variant proteogenomics from method development to biological applications.

## Methods

Figure 1A shows the overall workflow used in this study for proteogenomics searches against simulation, Alzheimer’s and HEK293 cell line data for variant identification and evaluation using PgxSAVy tool. Figure 1B depicts the steps in the PgxSAVy analysis, beginning from data input, rescoring, VAS calculation, statistical assessment, isobaric evaluation and disease annotation.

### Development of PgxSAVy

We utilized various PSM and variant event features for segregating true and false variant peptides for the development of a variant ambiguity score (VAS). The variant evaluation features are grouped as: match quality, variant events and result features. The PgxSAVy tool evaluates variant PSMs individually and does not conflate them into variant peptides before evaluation, as different scans matching to the same variant sequence may have variable quality. Here, we summarize the main features (See Supplementary Note 1 for details).

Match features include fractional intensity coverage (FIC), b and y ions continuity count (byCC), complementary ions and variant rescoring.

FIC is defined as-

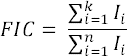

where,

n = total experimental peaks,

k = matched peaks

I_i_ = i^th^ matched peak intensity

b and y ions continuity count (byCC) is calculated as –

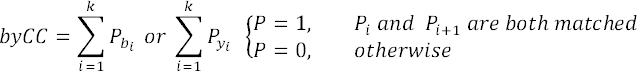

where,

k = matched peaks,

P_bi_ = b-ion flag (0 if absent, 1 if present),

P_yi_ = y-ion flag (0 if absent, 1 if present).

All the above features and complementary ions are used in MassWiz score (MW) [41]. So, we utilized this function to rescore the variant peptides for an additional layer of confidence. However, as it was shown to have a slight bias for longer peptides [42], we normalized the score by dividing it with the theoretical b & y ion counts. The normalized MassWiz Score (nMW) thus obtained, is calculated as: -

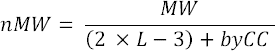

where,

L = Peptide length (2L-3 denotes theoretical b/y ions and assumes b1 to be absent),

byCC = b and y ions continuity count (explained before).

This normalization ensures that the nMW represents the strength of match per ion matched, and better intensity matches are still scored higher.

The variant event needs to be correctly localized at the appropriate amino acid, a strategy used for localizing post-translational modifications [43–45]. The best-scoring shuffled-decoy variant should score less than the true variant peptide reflected in its score difference ΔSV[34]. The score difference with the corresponding wild type (WT) peptide score should ideally be lesser (ΔWT).

The number of search engines identifying the variant peptide and the number of PSMs identifying a variant are also considered important features. Higher the number of search engines that identify a given MS/MS spectrum with the same variant peptide sequence, higher the probability of the PSM to be correct. Similarly, higher the number of PSMs identifying a variant sequence, higher are its chances of being correct.

The Variant Ambiguity Score (VAS), calculates a score that can assess the quality of the variant peptides which are classified as– confident, semi-confident or doubtful. VAS for a variant *i*, is defined as –

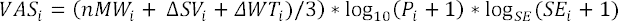

where,

nMW_i_ = normalized MassWiz Score for i^th^ variant match,

ΔSV_i_ = delta shuffled variant score for i^th^ variant match,

ΔWT_i_ = delta wildtype score for i^th^ variant match,

P_i_ = PSM counts for i^th^ variant match,

SE = number of search engines used for the proteogenomics search (this remains constant for a study and is used as the logarithmic base for calculating SE_i_ weight),

SE_i_ = number of search engines that identified the i^th^ variant PSM.

Isobaric (or near isobaric) amino acids and modifications are indistinguishable if these masses are within the instrument mass error [46]. We implemented an isobaric assessment module, that evaluates such mass difference and can result in three broad classes (i) single variant (ii) double variant, and (iii) isobaric. For the identified variants, PgxSAVy performs disease annotation using biological information present in the UniProt database that includes clinical significance and phenotype. A Perl script was also written to create batch file for the PSMs for annotation and visualization through pLabel tool.

All the above steps are implemented in the tool PgxSAVy, that reads the input, rescores variants in an automated manner, calculates VAS scores and classifies the variants with isobaric checks and perform biological annotation. PgxSAVy is freely available for academic use at https://github.com/anuragraj/PgxSAVy.

### Datasets

#### Simulation data

A simulated data was generated using MaSS-Simulator [47] using various parameters: dissociation strategies, noise content, relative intensities of signal peaks, peptide coverage, peptide length and peptide modifications (Supplementary Table S1 and S2). The input provided to the MS/MS spectra simulator contained 5000 peptides with single amino acid variation and 5000 peptides with two amino acid variations along with 40,000 wildtype peptides containing no variation (Figure 2).

#### Alzheimer’s Disease (AD) data

A publicly available MS/MS data [48] from post mortem brain samples of Alzheimer disease patients was downloaded from PRIDE repository (PXD004010). The dataset had 1.7 million raw MS/MS spectra and converted to de facto data standard-“MGF” format using MSConvert tool from ProteoWizard [49].

#### HEK293 cell line data (HEK)

This dataset generated from HEK293 human embryonic kidney cell lines, was downloaded from pride repository (PXD001468) [50]. The dataset contained 1.1 million MS/MS spectra acquired on Q-Exactive orbitrap Mass spectrometer to evaluate wide precursor mass tolerance search for discovery of modifications. The data was also converted to MGF format with the help of MSConvert tool.

### Database Creation

#### Simulation database (SimDB)

Simulated data was search against SimDB containing 5,000 single and 5,000 double amino acid variant peptides with 500,000 unrelated (UR) variant peptides. These unrelated variant peptides had no overlap with the variant peptides used in simulation. The sequences for 40,000 wildtype peptides for which spectra were generated, were not added to search database.

#### Reference database (RefDB)

Brain AD dataset and HEK293 cell line dataset were searched in three stages. Reference database (RefDB) downloaded from SwissProt (May 2022) contained curated 42,362 protein sequences (proteins+isoforms).

#### GENCODE database (TranscriptDB)

We downloaded the human GENCODE database (version 40) [51] containing 246,624 transcripts were translated in three-frames, resulting in 1,676,122 ORFs using an in-house Perl Script.

#### Variant database (VariantDB)

For creating variant database (VariantDB), human protein and variant sequences from neXtProt database were downloaded in PEFF format [52]. Variant tryptic peptides were created, allowing a maximum of two variant amino acid residues per peptide. The VariantDB thus created contained 93,669,236 variant peptides.

To each of these databases, contaminant proteins downloaded from cRAP website (https://www.thegpm.org/crap/) were added to the FASTA before search.

### Database Search

#### Simulation data search

Simulation data was searched against SimDB using multiple database search engines (X!Tandem, OMSSA and MSGF+) with EuGenoSuite proteogenomics suite on a computer with 64GB RAM using 24 processing cores [19]. The search parameters used were: 0.1 Da precursor mass tolerance, trypsin as cleavage enzyme, no missed cleavage and carbamidomethylation at cysteine as fixed modification. The FDR cut-off was set at 1% using FDRScore method [53].

#### Proteogenomics Search exclusive with MSGF+ (single search engine)

One fraction from AD data (ad_pl01, hereafter referred to as F1) was searched with MSGF+ search engine (version v2019.07.03). Three searches were performed in stage-wise manner, in which every subsequent search was conducted on the unassigned spectra from previous search. The searches were conducted iteratively in ReferenceDB, TranscriptDB, and VariantDB. The parameters for the first and second search were: – 6ppm precursor mass tolerance, 2 missed cleavages and trypsin as the proteolytic enzyme, fixed modification of carbamidomethylation at cysteine and variable modification of methionine oxidation were used. For third search, no missed cleavage was allowed, keeping other parameters same. The FDR cut-off was set at 1% at every stage.

#### AD data search with multiple search engines

All 10 fractions (F1-F10, as per nomenclature used in this study) from the AD dataset were searched with EuGenoSuite [19] search pipeline in a stage-wise manner against ReferenceDB, TranscriptDB, and VariantDB. EuGenoSuite pipeline was used for this purpose which had the three search engines mentioned above. The first search was conducted on ReferenceDB, second stage on the unassigned spectra against TranscriptDB, and the third search on still unassigned spectra against VariantDB. The parameters for this first and second stage search were: – 6ppm precursor mass tolerance, 2 missed cleavages and trypsin as the proteolytic enzyme, fixed modification of carbamidomethylation at cysteine and variable modification of methionine oxidation were used. For third stage search, no missed cleavage was allowed, keeping other parameters same. The FDR cut-off was set at 1% at every stage.

#### HEK data search with multiple search engines

All fractions of HEK data were also searched in multi stage manner using EuGenoSuite in similar manner as described for AD dataset. The parameters for the search: – 5ppm parent mass tolerance, 0.01Da fragment mass tolerance, trypsin as the enzyme with one missed cleavage allowed. Carbamidomethylation at cysteine and oxidation at methionine were used as fixed and variable modifications respectively. The FDR cut-off was set to 1% at every stage.

### Global and Class-specific FDR estimation

For gFDR estimation, all PSMs from the three different search stages were combined together before FDR estimation using the combinedFDR method [53] for integrating multiple search algorithm results. For class-specific FDR (cFDR) estimation, the identified peptides were classified into – known, novel and variant peptides, and FDR estimated separately for each class. The three categories were created based on mapping the identified peptides on reference proteome using an in-house Perl script and labelling each peptide in one of these categories. The mismatches up to two amino acids are labelled as variants, and higher mismatches are labelled as novel peptides. For both gFDR and cFDR, the formula remained the same although the input is different:

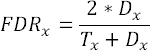

where,

*x* = category of FDR input (global i.e. all peptides, or class –i.e. known, novel or variants),

FDR_x_ = false discovery rate for particular category,

D_x_ = number of decoy PSMs above threshold for the category,

T_x_ = number of target PSMs above threshold for the category.

Based on the estimated FDR, 1% threshold was used and peptide hits passing the threshold are selected for further analysis. All variant PSMs were evaluated by PgxSAVy for quality assessment and annotation.

### Manual annotation of spectra for creating gold standard for VAS evaluation

We performed rigorous manual annotation of variant PSMs identified in the variant proteogenomic search conducted on F1 fraction of AD through multiple search engines. For all variant PSMs above 1% FDR, the four authors independently annotated each PSM as “good” or “bad”. Majority-voted “good” and “bad” PSM annotations were accepted, while the rest (ties) were labelled as “average”. The annotated PSM images for the spectra were generated through PLabel [54] using batch mode. A total of 797 PSMs were analysed and categorised in these three categories.

### Variant evaluation through PgxSAVy

All the variants in different searches were independently analyzed through PgxSAVy. For the PgxSAVy input, the EuGenoSuite search result was provided in tab-separated format with these parameters: search engine count, fragment tolerance, fixed modification type, variable modification type, instrument type, peak threshold, and number of minimum peaks. For single search engines, value of 1 was used.

### STRING analysis

For STRING analysis (version 12) [55], the proteins with all variants were compared against proteins with reliable variants (confident and semi-confident). The networks were created with high confidence threshold of 0.9 with only input proteins.

## Results and Discussion

### PgxSAVy Workflow

The overview of current study is shown in Figure 1A. Three different datasets from simulation, AD and HEK were used in this study to evaluate variants through PgxSAVy tool, the workflow for which is explained in Figure 1B. In PgxSAVy tool, we have integrated variant peptide rescoring, scoring variant PSM quality through VAS, isobaric assessment and disease related annotation of the variants. We tested the method using multiple datasets of increasing complexity. We used a simulation dataset, another dataset with manually annotated variant quality, and large-scale datasets for cataloguing the variant quality. The identified variant peptides from these datasets at 1% FDR were re-evaluated through PgxSAVy.

The method was first benchmarked on simulation data, which represents ideal scenario with near perfect separation of good and bad PSMs. But, since the simulated data differs from real-life spectra, we also used publicly available mass spectrometry data to evaluate the scoring method. The databases searches were conducted in multi-stage manner using a modified version of EuGenoSuite [19] which is a multi-algorithm proteogenomic search pipeline, and integrated ProteoStats to calculate false discovery rate (FDR) [56]. If the spectra from known and novel proteins are assigned at stage one and two respectively, these are removed from variant stage search and thus, it reduces the chances of random matching for variant peptides.

The combined FDR method [53] was used to estimate the integrated FDR from multi-search algorithms at 1% global FDR threshold (gFDR). In addition, a class-specific FDR approach (cFDR) was calculated for specific peptide classes i.e., known, novel and variant peptides after mapping the identified hits on the reference proteome. Eventually, the results were analyzed through PgxSAVy. The z-score is calculated from poor-quality hits that follow a normal distribution centred at zero (Supplementary Figure S1). The minimum value of incorrect scores of VAS (minVAS) is used to mirror and determine the maximum VAS for incorrect variants, as its absolute value |minVAS|. The null hypothesis H_0_, assumes that the scores following this null distribution are false. If a variant scores significantly higher, it is likely to be correct and incorrect otherwise. Using the z-score, p-value is calculated and variant PSMs are classified as (i) confident (p-value <= 0.01), (ii) semi-confident (p-value <=0.1) and (iii) doubtful (p-value >0.1).

**Figure 1:**
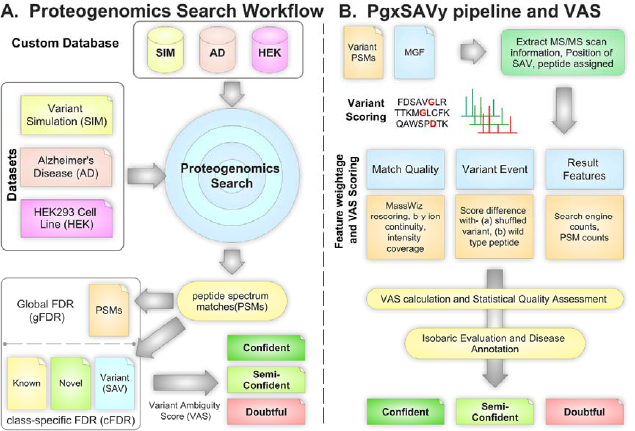
Workflow for identification and post-search validation of variant peptides from proteogenomics searches using PgxSAVy. (A) Simulation (SIM), Alzheimer’s disease (AD) and HEK293 Cell line (HEK) data were searched against respective databases-simDB, AD and HEK search database using EuGenoSuite, followed by global and class-specific (gFDR and cFDR) FDR estimation. Variant Ambiguity Score (VAS) is calculated for the variant PSMs that pass 1%FDR threshold through PgxSAVy. (B) In PgxSAVy tool, VAS is calculated using variant search results and MS/MS spectra, utilizing match quality, variant event and result features. This is followed by statistical quality assessment (z-score and p-value calculation) for quality evaluation and classification. Isobaric assessment and annotation is also performed within PgxSAVy framework for the variant PSMs.

We have implemented three types of annotation in PgxSAVy for variant peptides: (i) quality annotation (described above), (ii) variant class annotation (single variant, double variant, or isobaric), and (iii) disease annotation. Quality annotation includes assigning variant peptide quality such as – confident, semi-confident and doubtful based on p-values calculated from VAS. These annotations help in determining about the quality of variant peptides and whether it should be included in downstream analysis. The confident, and semi-confident variants are considered true variants and doubtful are considered as false. Isobaric analysis is used to annotate the variants as single amino acid variant, double amino acid variant, isobaric or modification peptide or a combination of these. If any variant amino acid in a peptide has a mass shift which can be justified with post translational modification shift, the variant peptide should not be considered as a variant amino acid. For peptides with two amino acid variants, it can be classified based on whether both amino acids are variants, or a single variant with isobaric or modification mass (Supplementary Figure S2). The disease annotation includes annotation of these variant peptides with biological information present in the UniProt database about its clinical significance and phenotype.

### Evaluation of VAS on simulated data

Simulated data for 10,000 variant peptides containing one or two variants per peptide sequence (5000 each) was generated using MaSS-Simulator as explained in Supplementary Table S1 and Figure 2A. Additionally, spectra were simulated for 40,000 wild type peptides, which were not-related to the variant sequences used above. All 50,000 spectra were generated with 20% noise to simulate non-perfect matching. The database for searching this data contained all sequences except the 40,000 WT peptides which can cause some WT peptides to match variants (false positives). We also added 5,00,000 variant peptides in database for which no spectra were generated and any spectra matching to these sequences will also be false. This enabled us to rigorously test the PgxSAVy pipeline with known true and false hits. The data was searched using EuGenoSuite applying three search algorithms as describe in methods. Variant PSMs obtained at 1% FDR were evaluated with PgxSAVy.

The density plots of various features used in PgxSAVy and their discriminatory power simulation results is shown in Supplementary Figure S3, and the density distribution of correct and incorrect variants show good separation by VAS (Supplementary Figure S4). After removing decoy and contaminants from 1% FDR results, a total of 11326 PSMs remained. The true positives and false positives in the results were 9905 (87.45%) and 1421 (12.54%) respectively. The search failed to identify 95 (0.95%) variants which were false negatives. This highlights that 1421 false positive variants (12.54%) could creep in the search results of near-ideal PSMs in simulation data. This also suggests that real world data being less than ideal, might have much more false variant identifications. Evaluating the quality of 11326 variant PSMs with PgxSAVy, classified 9775 variant PSMs as true (4731 confident and 5044 semi-confident), while the remaining 1551 PSMs as doubtful (Figure 2B). From these 9775 PSMs, only 154 (1.57%) incorrect PSMs were misjudged as either confident or semi-confident, while 9621 (98.43%) were correctly identified as true. The true identification percentage in final results improved from 87.45% to 98.43% which shows that PgxSAVy enhances the true identification rates. Furthermore, only 284 (1.83%) of correct PSMs were mis-classified as doubtful. Also, 1267 (89.16%) false variants were correctly labeled and thus removed. This shows PgxSAVy was able to effectively evaluate majority of true as well as false variants correctly, and performs with high sensitivity and specificity. PgxSAVy also analyzed the isobaric type classification for the true variant PSMs and found 4994 and 4584 were correctly annotated from the simulated single and double variants (5000 each) respectively. Of this, 4635 (92.81%) and 4584 (91.68%) were correctly classified as single and double variants. A small number of variant PSMs were mis-annotated from double to single variant class or in isobaric class (Figure 2C).

**Figure 2:**
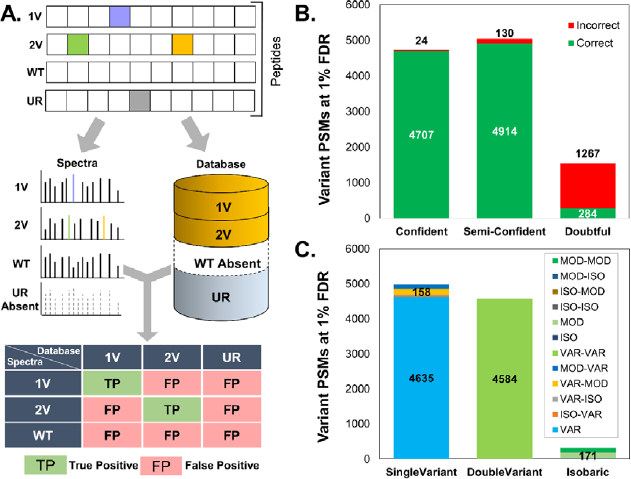
(A) Workflow for generation of simulated spectra and the database construction with true and false positive evaluation matrix. Simulated spectra (5000 each) were created for single variants (1V) and double variants (2V), along with 40000 wild type peptides (WT). 500,000 unrelated variants (UR) were added to FASTA database but spectra were not simulated for these. The 1V and 2V peptides were included in FASTA but not WT peptides. Based on this, the evaluation table shows the definition of true and false variant peptides. (B) The proportion of correct and incorrect variants is shown in the PgxSAVy classified variants. (C) Proportion of single, double and isobaric PSMs after isobaric evaluation by PgxSAVy.

### Evaluation on the manually validated dataset

The simulated dataset does not entirely reflect the real world scenario in terms of deviations induced due to sample complexity, technical variations, noise etc. However, it is a challenge to evaluate pan proteogenomics methods on real-world data, due to lack of well-established ground truth data – i.e. data with well-known true or false variant PSMs. For this purpose, we manually annotated the variant spectra as *Good*, *Average* or *Bad* quality (see methods) from F1 fraction of the AD data to establish ground truth for evaluation purposes. The F1 data contains 172,337 spectra on which a proteogenomics search was conducted using EuGenoSuite, followed by variant evaluation by PgxSAVy into *Confident*, *Semi-Confident* and *Doubtful* PSMs.

The proteogenomics search led to identification of 60589 known, 185 novel and 797 variant PSMs, of which we only selected the 797 variant PSMs for further analysis. Through manual validation, 221, 363 and 213 PSMs were categorized in the good, average and bad category, respectively. The density plots and scatter plots of various features against VAS is shown with their manual annotation class in Supplementary Figure S5 & S6 respectively.

Out of 797 variant PSMs, 270, 99 and 428 were identified as confident, semi-confident and doubtful PSMs respectively. We compared the density distribution of manual annotation with PgxSAVy quality annotations and found that their VAS distributions are highly similar (Figure 3A & B). This was also observed by comparing the means of the three classes in boxplots (Figure 3C & D) which were markedly similar despite some disagreements in the number of variant PSMs in each class-wise comparison (Supplementary Table S3). We observed that VAS was slightly stricter than manual annotation (especially for average quality PSMs). Thus, it is possible that some average labeled PSMs were either classified as confident or as doubtful by VAS. However, the overall mean VAS for each class remains similar (Figure 3C & D) which again shows that PgxSAVy is highly effective in variant quality estimation.

**Figure 3.**
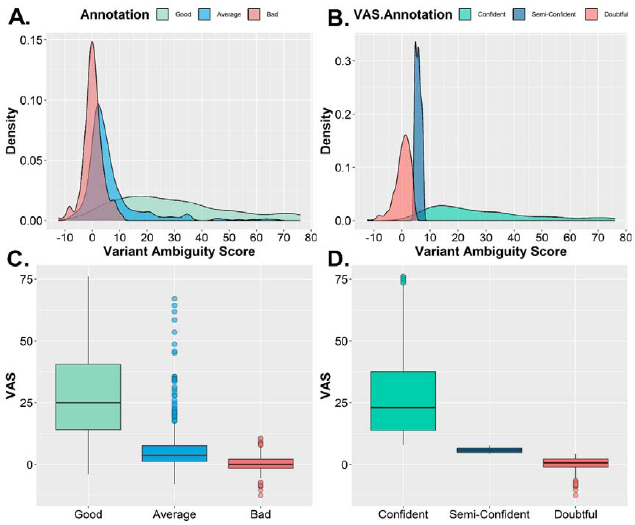
(A) Distribution of all variant PSMs identified in brain AD dataset (first fraction) with manually annotation (B) Distribution of all variant PSMs identified in brain AD dataset (first fraction) with VAS quality annotation.

### Can multi-algorithm searches and cFDR remove false variants?

We have demonstrated the effectiveness of PgxSAVy using simulated data and manually annotated data and found it to be highly effective in quality control of variant PSMs. For manually annotated evaluation, we had selected search results at 1% class-specific FDR conducted with multiple search engines. We had chosen this approach based on recommendations from previous variant proteogenomic studies [35,36]. Alfaro et al. suggested employing multiple algorithms, the cFDR method, and various post-search filters help us to identify confident variant peptides [36]. However, these aspects have not been systematically evaluated. Here, we aimed to evaluate these approaches to control variant false positives by using (i) multiple algorithms, (ii) cFDR, and (iii) multiple algorithms with cFDR. We also intended to compare these results with PgxSAVy for finding the most suitable approach.

We sought to compare the degree of false removal in single vs multiple algorithm search, as well as gFDR vs cFDR, using F1 data. The F1 fraction of AD data was searched with the MSGF+ Search algorithm and EuGenoSuite (for multiple search engines) in a three-stage search with gFDR and cFDR estimation for evaluating the number of false positives. The complete results obtained are provided in Table 1. We identified 1710 variant PSMs from MSGF+ search, of which 287 are classified as confident, 239 as semi-confident and 1184 as doubtful. From multiple algorithms, we obtained 728 variant PSMs, of which 232 are classified as confident, 98 as semi-confident and 398 as doubtful (Figure 4A & Table 1). While a single algorithm appears to identify more variants, but the extra 786 PSMs are false. Multiple algorithm results are integrated using the combinedFDR method, which helps in the removal of many false positives. It is also important to note that neither method is able to completely remove false variants when we use global FDR. From this data, we recommend that if resources allow, multiple algorithms searches should be conducted. We also asked if cFDR method can improve these results. Applying cFDR to MSGF+ search results identified 847 variant PSMs, of which 246 were classified as confident, 95 as semi-confident and 473 as doubtful by PgxSAVy (Figure 4B). This shows that cFDR performs better than gFDR for single search engine. The figure also shows that while cFDR with multiple search engines increases false variants slightly, the overall gain in correct variants is higher than gFDR and PgxSAVy can remove the false hits. The detailed analysis of gFDR and cFDR on all files from AD and HEK data are provided in supplementary figure S7 and supplementary tables S5 and S6. These results demonstrate that multiple algorithms and cFDR combination is more potent in reducing false variants while keeping the sensitivity high. We recommend that this combination is best for variant identification. If one cannot perform multi algorithm searches, cFDR is still recommended for keeping false variants to a low proportion.

**Table 1.** Single and multiple search engine results with global and class specific FDR and VAS classification on variant PSMs.

|  |  | Known | Novel | Variant |  |  |  |
| --- | --- | --- | --- | --- | --- | --- | --- |
|  |  |  |  | Total | Confident | Semi-Confident | Doubtful |
| **Single Search Algorithm** | gFDR | 57962 | 286 | 1710 | 287 | 239 | 1184 |
|  | cFDR | 60743 | 243 | 847 | 246 | 95 | 473 |
| **Multiple Search Algorithm** | gFDR | 60634 | 53 | 728 | 232 | 98 | 398 |
|  | cFDR | 60589 | 185 | 797 | 270 | 99 | 428 |

To evaluate whether the VAS annotated PSMs also agree with manual annotations for multiple algorithms with gFDR and cFDR, these results were compared to each other in Venn diagram (Figure 4C). It was observed that all PSMs in gFDR were encompassed within cFDR set. Out of 69 variants exclusive to cFDR, the exclusive manual (15 good, 31 average and 23 bad) annotations depict that although 46 variant PSMs were identified as confident/semi-confident, it also allowed 23 doubtful PSMs to pass (Figure 4C). Even though multiple algorithms with cFDR perform better, even this combination is not fully capable of removing false variants. The figure 4D shows that similar VAS is scored for the three categories in manul annotation as well as VAS classification demonstrating that VAS achieves a performance similar to rigorous manual validation in automated manner. It should not escape notice that despite these FDR methods, the false variants are still looming large in data, and thus tools like PgxSAVy will be indispensable for variant proteogenomics.

Thus, we have established the utility of PgxSAVy through simulation and manual annotation data. We have also established that multiple search engine workflow combined with cFDR method is best suited for variant proteogenomics followed by PgxSAVy evaluation of the variants thus identified.

**Figure 4:**
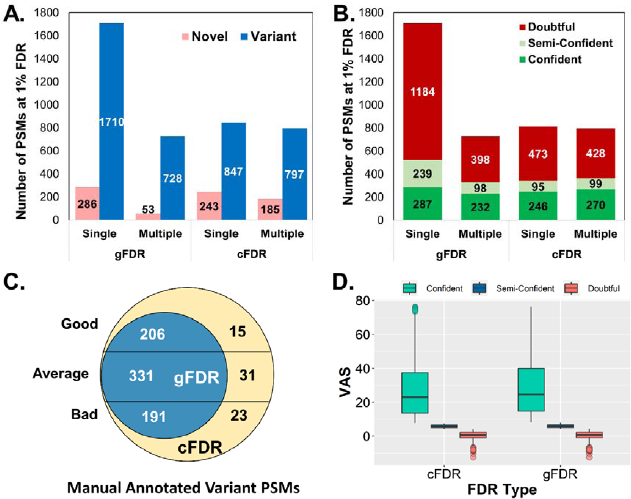
(A) Novel and Variant PSMs identification in single and multiple search engine results passing 1% FDR (B) PgxSAVy classification on variant PSMs showing higher ratio of classified confident PSMs with multiple search algorithm and cFDR (C) cFDR results encompass gFDR results and both methods have non-negligible number of PSMs categorized as bad by manual annotation. (D) VAS rescoring showing similar scores in for cFDR and gFDR for multiple search engine results but both methods are prone to false positives in variant PSMs which VAS can control better than the two methods of FDR.

### Analysis of variants in AD and HEK datasets through PgxSAVy

We applied PgxSAVy to two independent datasets, one from AD and another from HEK, performed variant quality control, and annotated the variants for their disease pathogenicity. This was performed to evaluate the effect of PgxSAVy on real world scenario with its implications on final biological outcome.

The AD dataset, containing 1,718,768 spectra from ten fractions, was searched using the EuGenoSuite. This dataset consisted of brain tissue in the Alzheimer’s disease condition from which a total of 595,892 known, 2671 novel, and 8883 variant PSMs were identified at 1% cFDR. After removing contaminants and decoy hits, the identified PSMs were 582986, 2671, and 8777 for the known, novel and variant PSMs respectively.

We selected the 8777 variant PSMs at 1% cFDR threshold for further analysis. PgxSAVy classified 2084 variant PSMs as confident, 987 as semi-confident and 5706 as doubtful (Table 2 and Figure 5A & B). From the reliable (confident and semi-confident) 3071 variant PSMs (1891 non-isobaric), we identified 782 unique variant peptides (500 non-isobaric) compared to 256 identified in another proteogenomics study on same dataset [40] (Figure 5C). Although we could not compare more details (information was not provided in the paper), we identified much higher number of true variants even after removal of doubtful and isobaric cases probably due to multiple algorithms and a robust statistical control. Another possible reason can be that the authors searched the data against sample-specific dataset, as compared to publicly available GENCODE reference database in this study. Searching in that database identifies a small fraction of variants (0.3%) that are sample-specific but a large proportion of variants from GENCODE database will not be identified, as also shown in an independent study [57].

**Table 2.** VAS classification for Brain AD dataset and HEK Cell line.

| VAS | AD dataset | HEK Cell line |
| --- | --- | --- |
| Total | 8777 | 3928 |
| Confident | 2083 | 945 |
| Semi-confident | 985 | 424 |
| Doubtful | 5709 | 2559 |

**Figure 5:**
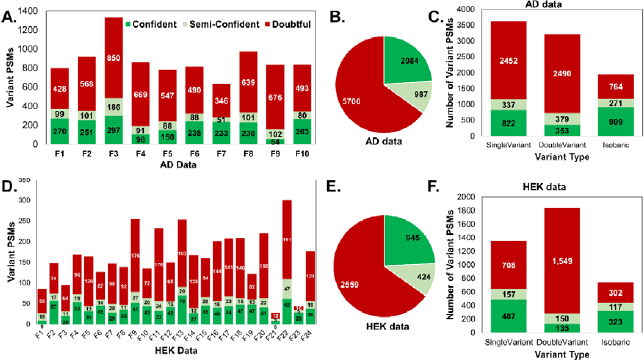
Distribution of identified PSMs (A) Variant PSMs classified by PgxSAVy for AD dataset for F1-F10 (B) Total Variant PSMs classified by PgxSAVy for AD data. (C) Isobaric analysis of reliable variants for AD data. (D) Variant PSMs classified by PgxSAVy for HEK dataset for F1-F24. (E) Total Variant PSMs classified by PgxSAVy for HEK data. (F) Isobaric analysis of reliable variants for AD data.

In the HEK data containing 1.12 million spectra, a total of 471,823 known, 1658 novel and 4015 variant PSMs were identified at 1% cFDR, of which 3928 remained after removing contaminants and decoys. PgxSAVy analysis classified 945 variant PSMs as confident, 424 as semi-confident and 2559 as doubtful (Table 2and Figure 5D & E). From the reliable (confident and semi-confident) 1369 PSMs (929 non-isobaric), we identified 626 unique variant peptides (434 non-isobaric) (Figure 5F). After removal of doubtful PSMs, 1159 single variant, 732 double variants and 1180 isobaric PSMs were left in AD data; while 644 single variant, 285 double variants and 440 isobaric PSMs remained in HEK data (Figure 5 C & F). The isobaric variant PSMs are those whose mass shifts can be explained by other amino acids or with common modifications. The PgxSAVy rescoring approach allowed for the successful detection of variant peptides in the HEK cell line data, enabling the investigation of their role in cellular processes and disease mechanisms.

The variants accepted after all filters were used to create the sunburst charts in Supplementary figure S8 and S9 that show the original amino acids and the frequency of conversion to the mutated amino acids in AD and HEK data respectively. Notably, in both independent datasets, even after several algorithms and strict cFDR, many false positives can sneak in, but PgxSAVy effectively removed those false variants, despite maintaining higher identification rate than previous study, which did not apply such strict filter. Approximately one-third of the variant peptides remained after stringent filtering by PgxSAVy and denotes its ability to identify the most reliable variant peptides. Furthermore, Supplementary Figure S10 displays the matched spectra of some identified variant peptides from different classes.

To investigate the biological significance of the findings, we analyzed the variants discovered in AD and HEK data in more detail. Figure 6 shows some variants identified in AD and HEK datasets. Neuromodulin protein (P17677-2) that plays a role in nerve growth was identified with mutations A214G (rs182766826) and A219V (rs577083604) in variant peptide QADVPAAVTAAGATTPVAEDAAAK supported with 20 reliable PSMs. Another protein Synapsin-2 (Q92777-2) was identified with mutation A127S (rs978216738) in variant peptide VLLVVDEPHADWSK (13 confident PSMs). Myelin proteolipid protein (P60201-2) was identified with one pathogenic mutation D168N (rs132630284) in variant peptide TSASIGSLCANAR (5 confident PSMs). In HEK data, a high-confidence peptide, TLNEADCAT[L/I]PPAIR with V609L/V609I mutation (rs6962) in succinate dehydrogenase (P31040-2) protein was identified with 88 confident PSMs and 3 semi-confident PSMs, which was also reported in original study [50]. For the same protein, we also found another variant peptide HTLSFVDVGTGK with Y581F mutation (rs6960) for the same protein which was not found earlier. Another new finding was Glucosidase 2 subunit beta (P14314-2), which has role in glycan metabolism, was also identified with variant peptide LGGSPTSLGTWSSWVGPDHDK having two mutations, G447S (rs761355123) and I450V (rs34351170). Ribosomal oxygenase 2 protein (Q8IUF8-4) was also identified with variant alanine-to-threonine, A385T (rs2172257) which plays an important role in cell growth and survival. We also discovered Triosephosphate isomerase (P60174-3) carrying a pathogenic mutation I208V, and another mutation G211S in peptide VVLAYEPVWAVGTSK with 8 reliable PSMs (7 confident, 1 semi-confident).

**Figure 6:**
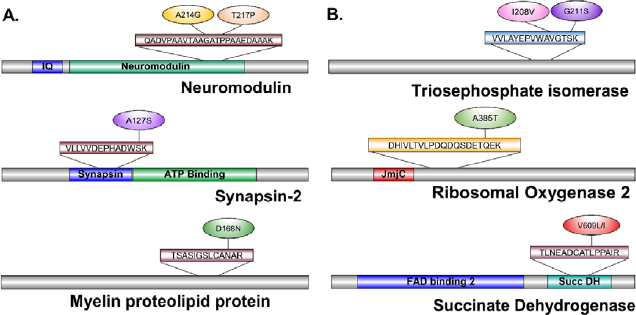
Some identified variants in (A)AD data (B) HEK data.

### Disease annotation of variants peptides in PgxSAVy

The quality control of variant peptides is not sufficient for establishing the functional implication of the identified variant peptides and thus, mapping information about their functional biological or disease impact is important. There are several rich sources of genomic variant assessment which need to be manually sifted for finding relevant information, a time consuming and tedious necessity. UniProt is a rich aggregated source of such information that we have leveraged towards an automated annotation of pathogenicity information for the identified variants.

PgxSAVy uses a downloaded annotation file from UniProt (https://ftp.uniprot.org/pub/databases/uniprot/current_release/knowledgebase/variants/homo_sapiens_variation.txt.gz) and maps the identified variant peptides for their annotations compiled in the UniProt database. This helps the biologists to make informed decisions about variant selection for further enquiry or planning validation experiments.

Using these annotations, we analyzed the important proteins using protein-protein interaction (PPI) database STRING. Here, we observed that while the PPI analysis of all proteins identifies all major pathways, PPI derived from a selective proteome set (only reliable variant peptides) leads to discovery of important pathways with mutated proteins. For example, glycolysis/gluconeogenesis mutations were highlighted in AD data; while spliceosome and actin folding in HEK data (Supplementary Figure S11 and S12).

## Conclusion

Detecting variant peptides through proteogenomics presents challenges in controlling false identifications, which cannot be tackled by cFDR. Various ad-hoc filters have limitations of scale. We developed PgxSAVy that performs an automated, comprehensive variant quality control, as well as annotates their biological implication. We demonstrated the utility of PgxSAVy on a simulated variant dataset and a manually validated dataset. We also showed that multiple algorithms perform better than single algorithms for curtailing the false variant effect, but cannot completely remove it. Furthermore, although the cFDR method provides better results than gFDR, but remains ineffective in reducing false variants. Applying PgxSAVy on large-scale public datasets, we show that a large number of variants are false and it is critical to employ quality control measure before interpretation. After a comprehensive scrutiny, we discovered several variants described in the original studies as well as discovering ones not described earlier.

In summary, our study underscores the effectiveness of PgxSAVy in controlling false positives in variant proteogenomics. We provide insights into its performance across different datasets and emphasize the biological significance of the findings. PgxSAVy is a handy tool that can perform post search variant quality control. It signifies the need for removing false variants that inadvertently use up limited critical resources for chasing incorrect findings. It can also help in biological evaluation of variants by proving facile integration with neXtProt knowledge for variant annotation and highlighting their role in disease, if any. Thus, PgxSAVy is a handy, comprehensive and automated tool for performing variant quality control and post-search-filters in variant proteogenomics.

## Data Availability Statement

All data are incorporated into the article and its online supplementary material.

## Figure legends

*Supplementary Figure S1: Density distribution of poor quality variant PSMs over VAS showing a normal distribution centred at zero for (A) simulation dataset and (B) manually annotated dataset*.

*Supplementary Figure S2: Flowchart for isobaric assessment of variant peptides in isobaric check module of PgxSAVy. The yellow nodes are steps or checks in the module, while the green nodes represent decision steps in the branch*.

*Supplementary Figure S3: Distribution of features for simulation data. Distribution of features for – match quality (A, B, C & D), variant events features (E, F), and result features (G, H). The true and false variants are shown in green and red respectively*.

*Supplementary Figure S4: Density distribution of VAS scores for (A) correct and incorrect variant peptides (B) VAS quality annotation, for simulation dataset*.

*Supplementary Figure S5: Distribution of features for manually annotated spectra to visually inspect their impact on separation of variant classes into good, average and bad quality*.

*Distribution of features for – match quality (A, B, C & D), variant events features (E, F), and result features (G, H). The good, average and bad PSMs are shown in green, blue and red respectively*.

*Supplementary Figure S6: Scatter plot for showing segregation of different features against VAS in AD dataset fraction F1, with manual annotations which shows good (green), average (blue) and bad (red) PSMs*.

*Supplementary Figure S7: Distribution of identified PSMs passing the 1% FDR threshold in gFDR and cFDR for novel and variant categories for (A) AD datasets (B) HEK293 cell line datasets*.

*Supplementary Figure S8: Sunburst diagram depicting the most common variant types found in AD data for the reliable variants after removing all isobaric PSMs*.

*Supplementary Figure S9: Sunburst diagram depicting the most common variant types found in HEK data for the reliable variants after removing all isobaric PSMs*.

*Supplementary Figure S10: Examples for spectra classified with VAS in (A) confident, (B) semi-confident, (C) doubtful category*.

*Supplementary Figure S11: AD all variant genes (left) depict PPI network of all pathways in non-specific manner. When selected high quality variants are used, glycolytic/gluconeogenesis pathways are highlighted*.

*Supplementary Figure S12: HEK all variant genes (left) depict PPI network of all pathways in non-specific manner. When selected high quality variants are used, spliceosomes and actin folding pathways by CCT/Tric are highlighted*.

## Table legends

*Supplementary Table S1: MaSS-Simulator Parameters used for generation of simulated spectra*

*Supplementary Table S2: Composition of peptides used for simulation spectra generation (MS/MS) and sequences in the search database. The strategy behind the decisions of this design with identified, missed, and falsely identified PSMs are also provided*.

*Supplementary Table S3: VAS classification against manual validation classes for F1 fraction of AD dataset*.

*Supplementary Table S4: PgxSAVy output on AD dataset comparing gFDR and cFDR. Supplementary Table S5: PgxSAVy output on HEK dataset comparing gFDR and cFDR*.

