## SupplementaryFiguresTables for "Quality control of variant peptides identified through proteogenomics- catching the (un)usual suspects"

**Supplementary Note 1: PgxSAVy development and features**

We utilized various PSM and variant event features for segregating true and false variant peptides for the development of a variant ambiguity score (VAS). Features like spectrum intensity coverage, b and y ion peak matches and their continuity score are well-known for evaluation of PSM quality and part of MassWiz score (MW). In addition, the continuity of a series (b/y) significantly enhances confidence in matched ions even when fragmentation is incomplete due to partially mobile or non-mobile proton-containing peptides. Since these features are generally not used together and also not part of search output, it makes the post-search evaluation on these features impossible. The variant evaluation features or descriptors can be grouped in three main categories: match quality, variant events and result features. These features have the ability to discriminate between good and bad variant hits. The PgxSAVy tool evaluates variant PSMs individually and does not conflate them into variant peptides before evaluation, as different scans matching to the same variant sequence may have a variable degree of quality.

**Match quality (Variant Rescoring)**

*Fractional intensity coverage (FIC)*

Fractional intensity coverage refers to the ratio of the summed intensity for the matched peaks for the peptide sequence, against total summed intensity of the respective spectrum.

$$FIC= \frac{\sum_{i=1}^{k} I_{i}}{\sum_{i=1}^{n} I_{i}}$$

where,

n = total experimental peaks,

k = matched peaks

I_i_ = i^th^ matched peak intensity

A peptide that covers most of the high-quality MS/MS peaks for a spectrum gets a good fractional intensity coverage. Thus, this feature can distinguish between good and bad PSM quite well and is part of the variant scoring.

*b and y ions continuity count (byCC)*

Occurrence of continuous b and y ions in a spectrum provides information about the quality of spectrum match and can distinguish a true variant from false one. More the number of continuous b and y ion ladders, more the indication of good fragmentation and quality variant match.

$$byCC=\sum_{i=1}^{k} P_{b_{i}} or \sum_{i=1}^{k} P_{y_{i}} \left\{ \begin{aligned} P=1, &P_{i} {and P}_{i+1} are both matched \\ P=0, &otherwise \end{aligned} \right.$$

where,

k = matched peaks,

P_bi_ = b-ion flag (0 if absent, 1 if present),

P_yi_ = y-ion flag (0 if absent, 1 if present).

*Complementary ions*

Complementary ions such as- (i) immonium ions, (ii) neutral losses of water, and (iii) neutral losses of ammonia, can also aid in distinguishing between borderline true and false hits when b and y ions counts are closely similar, which can immensely benefit the identification of variant peptides.

*Variant peptide rescoring*

All the above features are present in MassWiz algorithm (Yadav, Kumar and Dash, 2011b). The scoring system comprehensively utilizes the MS/MS information present in a spectrum, and provides an effective score for every PSM. So, we utilized this function to rescore the variant peptides for an additional layer of confidence. However, as it was shown to have a slight bias for longer peptides (Yadav, Kumar and Dash, 2012), we fixed this bias by normalizing it using the theoretical b & y ion counts. The normalized MassWiz Score (nMW) thus obtained, is calculated as: -

$$nMW= \frac{MW}{\left( 2 \times L-3 \right)+byCC}$$

where,

L = Peptide length (2L-3 denotes theoretical b/y ions and assumes b1 to be absent),

byCC = b and y ions continuity count (explained before).

This normalization ensures that the nMW represents the strength of match per ion matched, and better intensity matches are still scored higher.

**Variant Event Features**

*Score difference with shuffled variant decoy peptides (ΔSV)*

The variant event needs to be correctly localized at the appropriate amino acid to distinguish from other possible variant positions in the peptide sequence. Previously, post translational modifications studies have used this strategy to build a rescoring to correctly localize the modification site (Fermin *et al.*, 2013; Aggarwal *et al.*, 2021, 2023). The hypothesis is that a decoy variant peptide (created by swapping the variant amino acid with other amino acid, one at a time) in the sequence, should always score less than a correct variant peptide. This ΔSV score highlights the gap between a correct variant and a random decoy variants, as exemplified in a previous study (Yi *et al.*, 2018).

*Score difference with wild type peptide (ΔWT)*

For the correct variant peptide match, the corresponding score for the wild type (WT) peptide should ideally be lesser. Higher the score difference between variant peptide and WT peptide, higher the chances of variant match to be correct.

**Result features**

*Number of search engines identifying the variant peptide*

Higher the number of search engines that identify a given MS/MS spectrum with the same variant peptide sequence, higher the probability of the PSM to be correct. During a database search, it is plausible that different search engines report different peptide sequences for the same scan. For variants, search engines are more likely to agree for true variants, while disagreements may be due to: - (i) false hits, (ii) low information content in MS/MS, or (iii) one search engine reporting incorrect variant. Thus, using search engine count as a weightage can add additional discriminatory power.

*PSM count per peptide*

For any identified variant, higher the number of PSMs identifying the same variant sequence, higher are the chances of the variant being correct. A majority of incorrect hits are identified with a single PSM which requires greater scrutiny. The chances of random matching of variant peptides to single PSMs in proteogenomics is very high, and such “one-hit proteogenomics-wonders” occur at higher frequency than in proteomics. So, PSM counts per variant peptide can prove to be a useful indicator for correct variants.

**Variant Ambiguity Score (VAS)**

The VAS calculates a score that can assess the quality of the variant peptides which are divided into three classes – confident, semi-confident and doubtful. For a variant peptide spectrum match i, the VAS is defined as –

$${VAS}_{i}=\left( {nMW}_{i}+ {\Delta SV}_{i}+{\Delta WT}_{i})/3 \right)*\log_{10} \left( P_{i}+1 \right)*\log_{SE} ({SE}_{i}+1)$$

(eq 1)

where,

nMW_i_ = normalized MassWiz Score for i^th^ variant match,

ΔSV_i_ = delta shuffled variant score for i^th^ variant match,

ΔWT_i_ = delta wildtype score for i^th^ variant match,

P_i_ = PSM counts for i^th^ variant match,

SE = number of search engines used for the proteogenomics search (this remains constant for a study and is used as the logarithmic base for calculating SE_i_ weight),

SE_i_ = number of search engines that identified the i^th^ variant PSM.

A Perl script was also written to create batch file for the PSMs for annotation and visualization through pLabel tool.

*Isobaric analysis on identified variant peptides through proteogenomics*

Isobaric (or near isobaric) amino acids and modifications are indistinguishable if these masses are within the instrument mass error (Deutsch *et al.*, 2019). To tag such peptides, we implemented an isobaric assessment module, that evaluates the mass difference between variant amino acids or modifications within the applied MS/MS tolerance range. This check is performed for both single and double variant types and can result in three broad classes and their respective subclasses (in parentheses)– (i) single variant (var, iso-var, mod-var), (ii) double variant (var-var), and (iii) isobaric (iso, iso-iso, iso-mod).

*Annotating identified variants through PgxSAVy*

For the identified variants, PgxSAVy performs disease annotation using biological information present in the UniProt database that describes their pathogenicity. These biological annotations include their clinical significance and phenotypes. If the annotation is missing in curated databases culled by UniProt, it could be a novel variant discovery (not observed before).

All the above steps are implemented in the tool PgxSAVy, that reads the input, rescores variants in an automated manner, calculates VAS scores and classifies the variants with isobaric checks and perform biological annotation.

**Supplementary Figures**

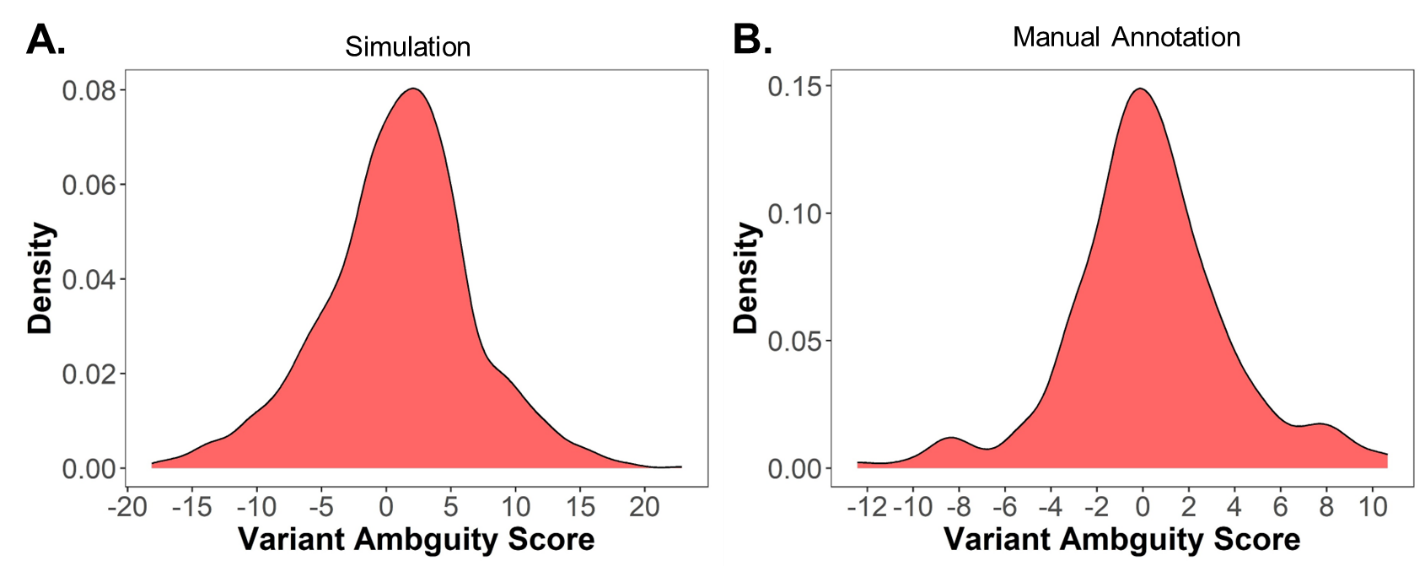

*Supplementary Figure S1: Density distribution of poor quality variant PSMs over VAS showing a normal distribution centred at zero for (A) simulation dataset and (B) manually annotated dataset.*

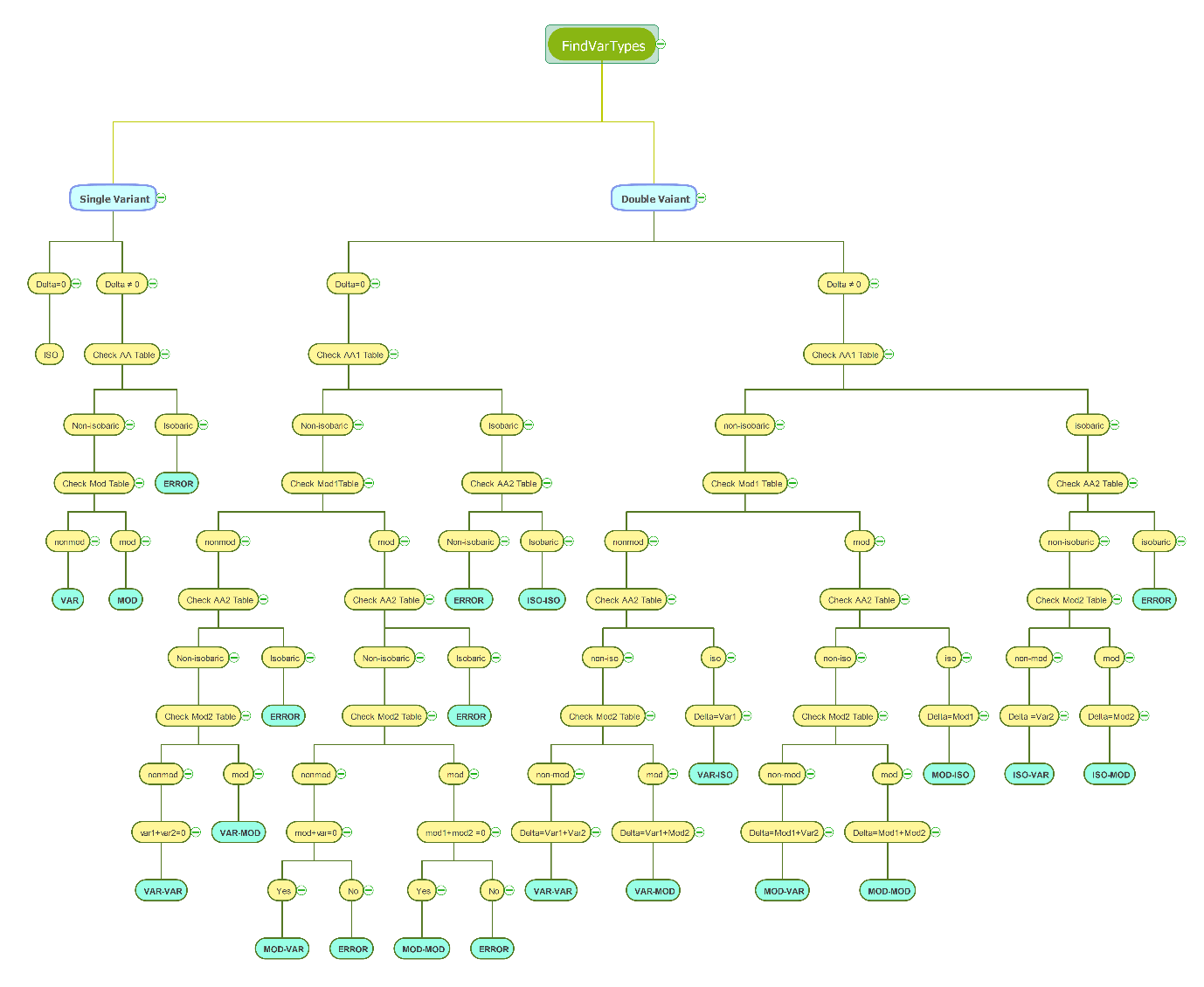

*Supplementary Figure S2: Flowchart for isobaric assessment of variant peptides in isobaric check module of PgxSAVy. The yellow nodes are steps or checks in the module, while the green nodes represent decision steps in the branch.*

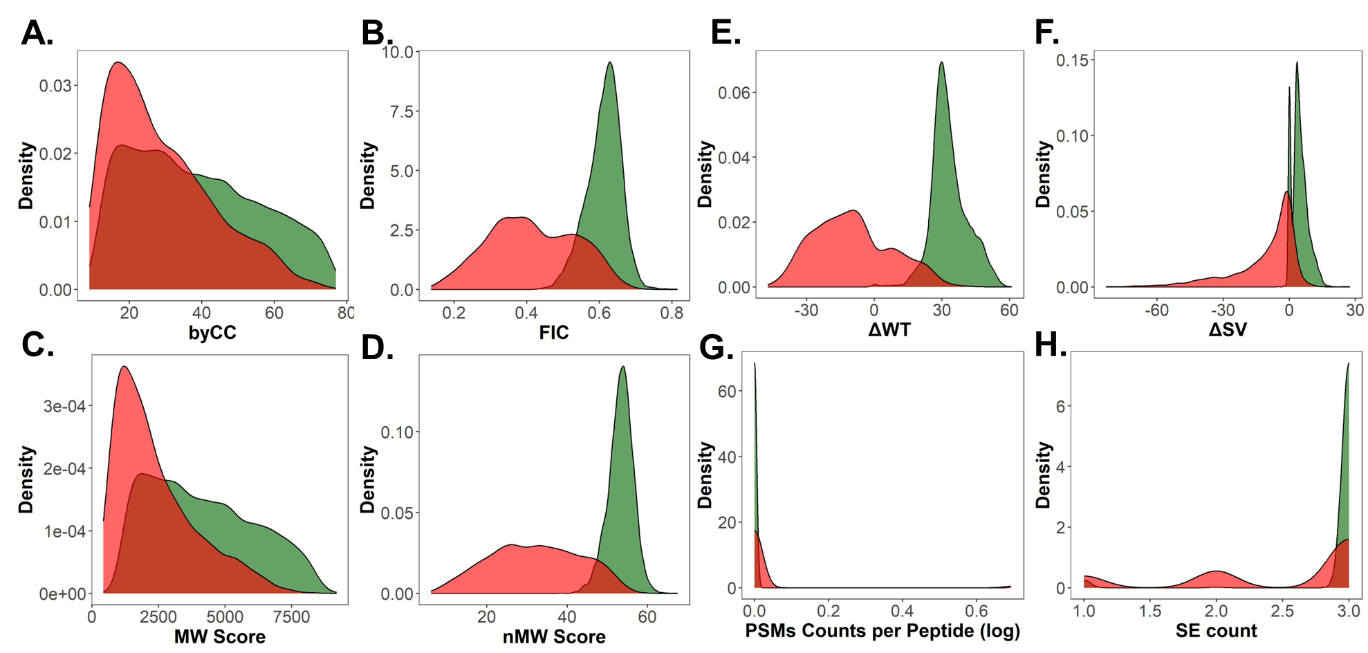

*Supplementary Figure S3: Distribution of features for simulation data. Distribution of features for – match quality (A, B, C & D), variant events features (E, F), and result features (G, H). The true and false variants are shown in green and red respectively.*

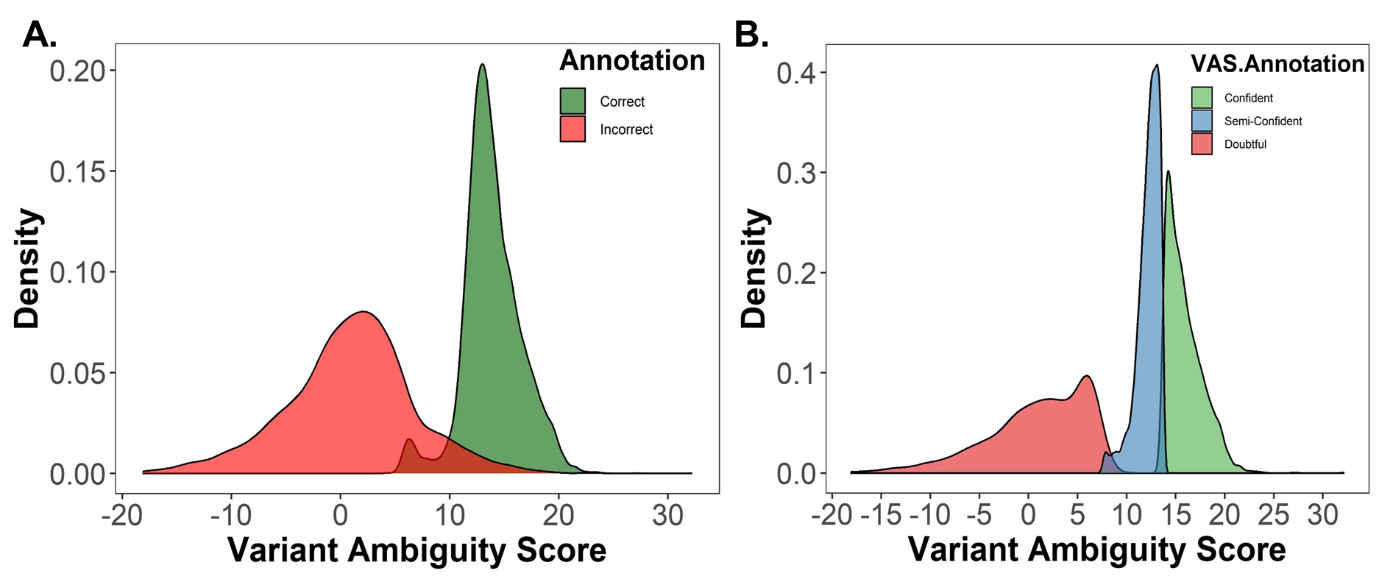

*Supplementary Figure S4: Density distribution of VAS scores for (A) correct and incorrect variant peptides (B) VAS quality annotation, for simulation dataset.*

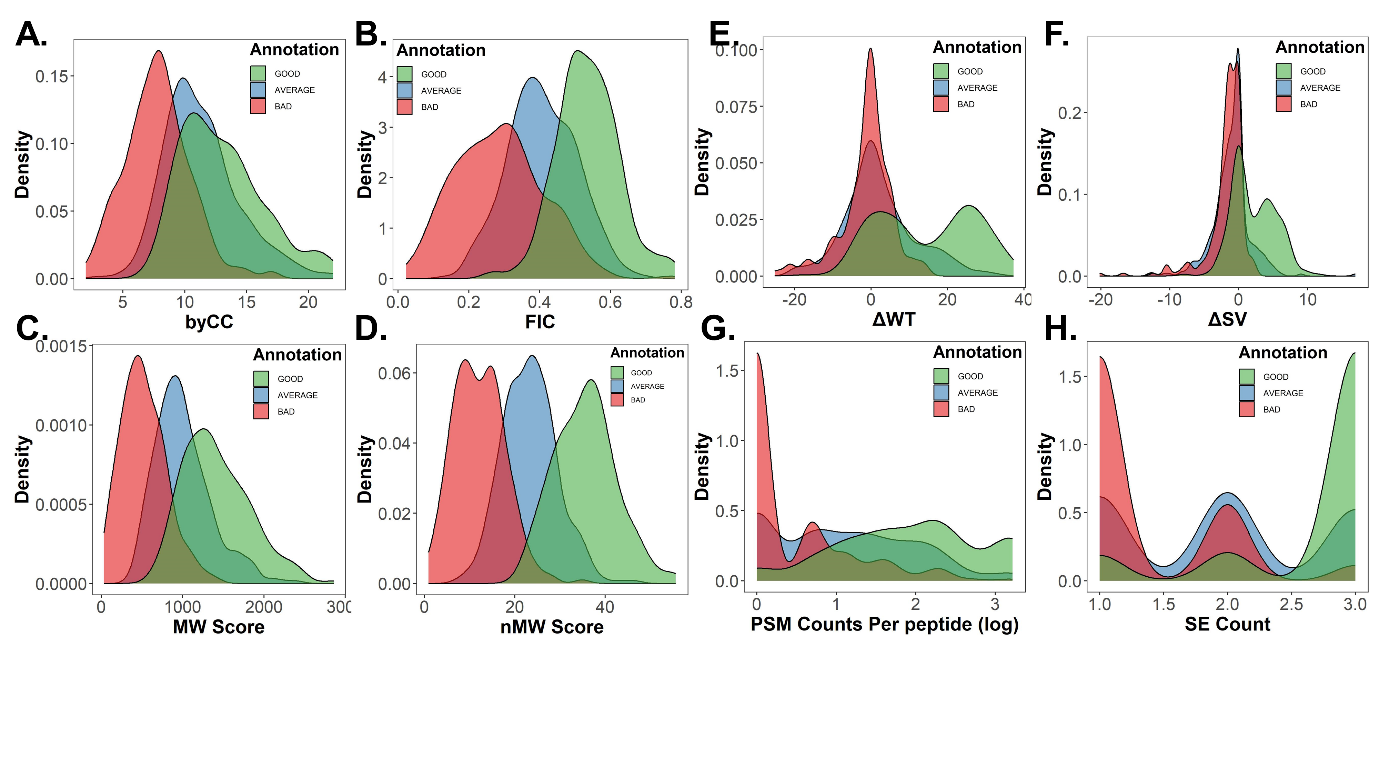

*Supplementary Figure S5: Distribution of features for manually annotated spectra to visually inspect their impact on separation of variant classes into good, average and bad quality. Distribution of features for – match quality (A, B, C & D), variant events features (E, F), and result features (G, H). The good, average and bad PSMs are shown in green, blue and red respectively.*

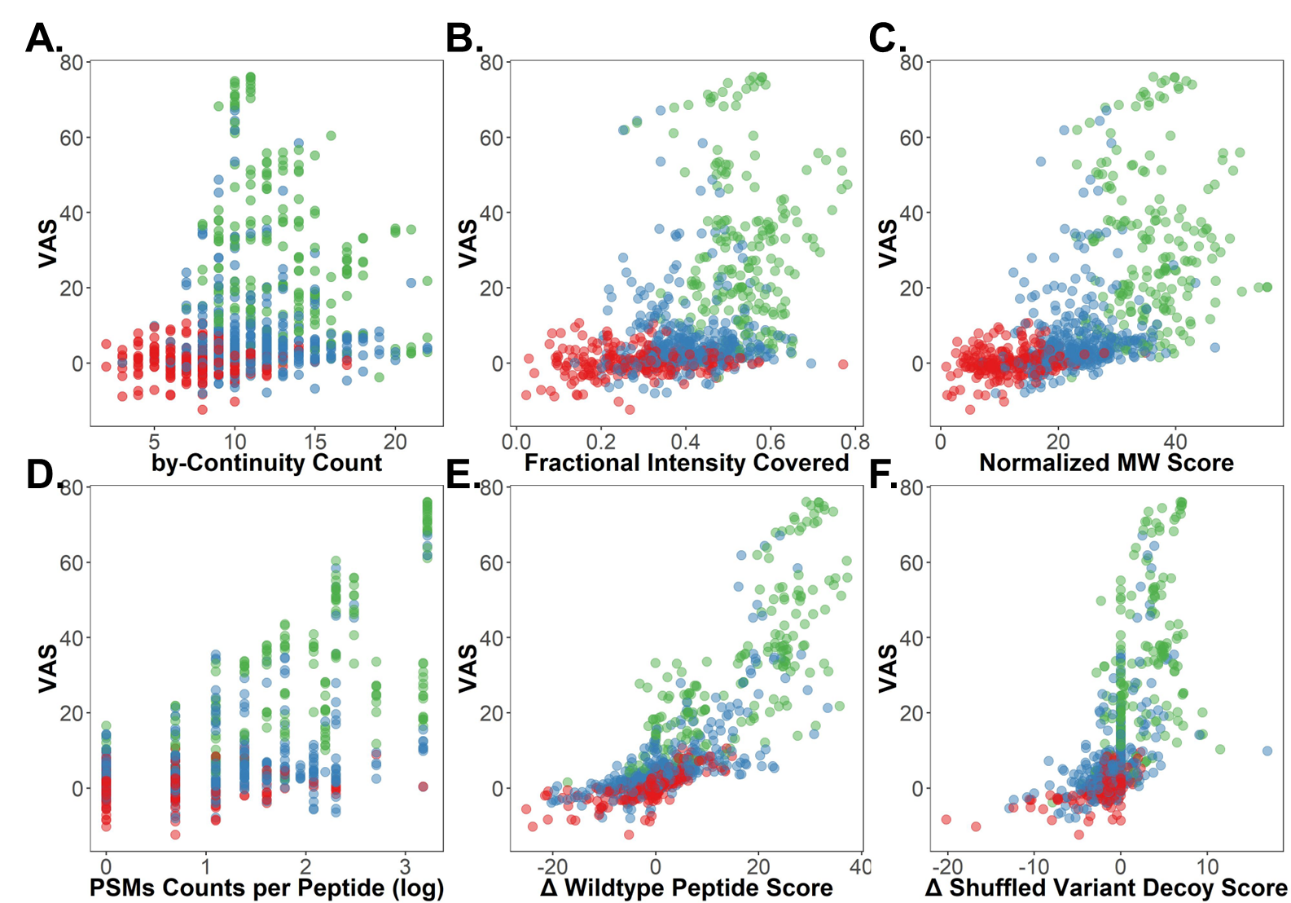

*Supplementary Figure S6: Scatter plot for showing segregation of different features against VAS in AD dataset fraction F1, with manual annotations which shows good (green), average (blue) and bad (red) PSMs.*

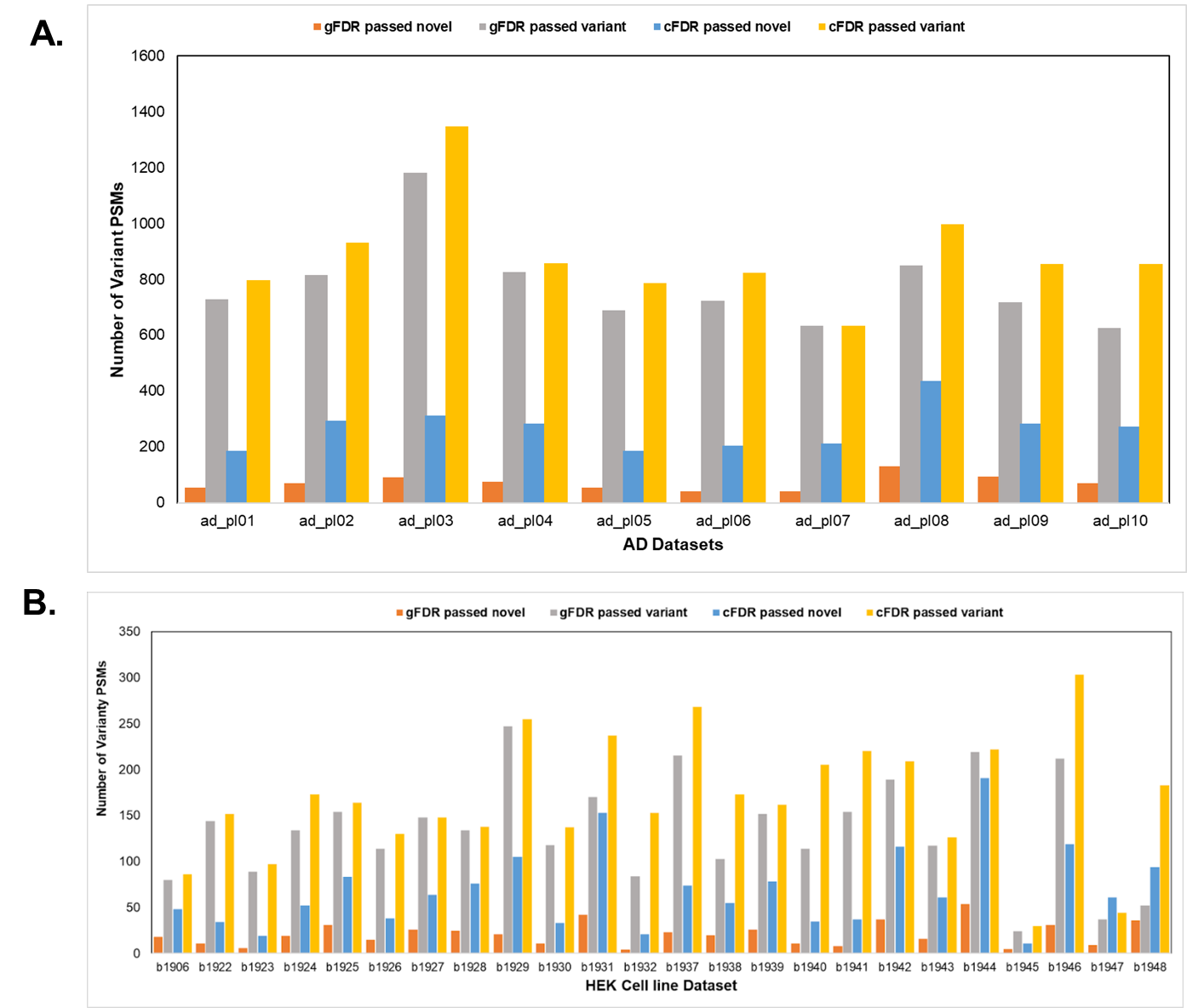

*Supplementary Figure S7: Distribution of identified PSMs passing the 1% FDR threshold in gFDR and cFDR for novel and variant categories for (A) AD datasets (B) HEK293 cell line datasets.*

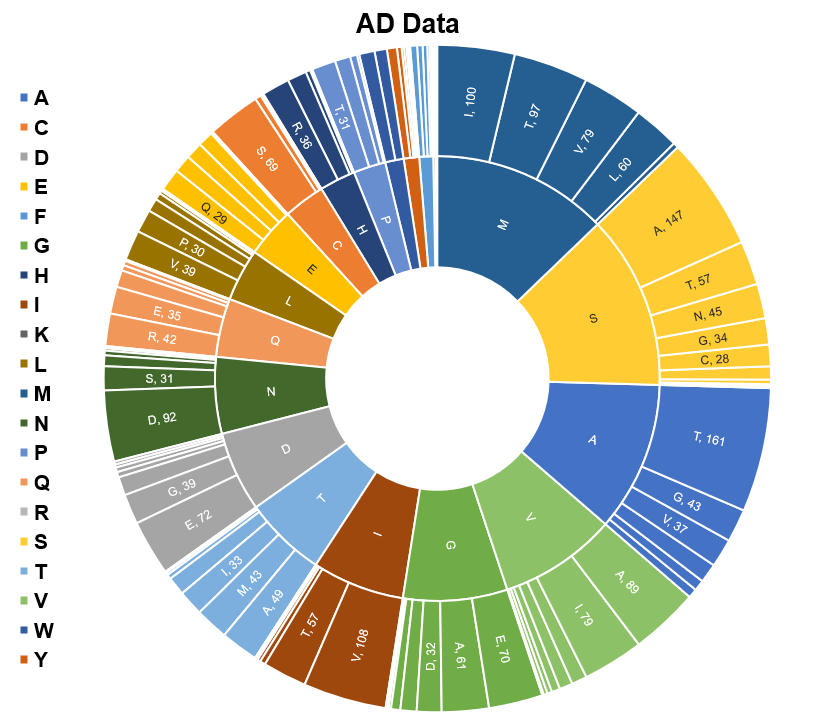
*Supplementary Figure S8: Sunburst diagram depicting the most common variant types found in AD data for the reliable variants after removing all isobaric PSMs.*

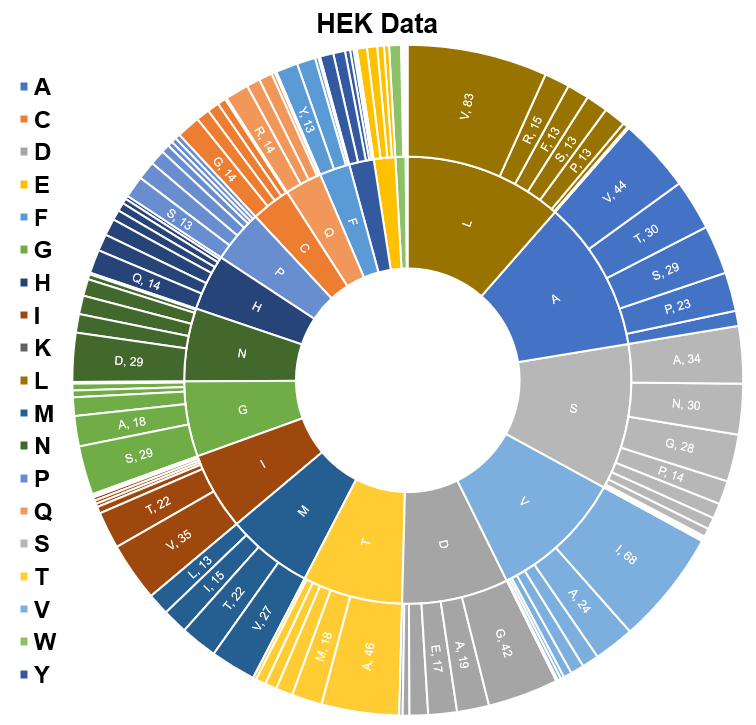
*Supplementary Figure S9: Sunburst diagram depicting the most common variant types found in HEK data for the reliable variants after removing all isobaric PSMs.*

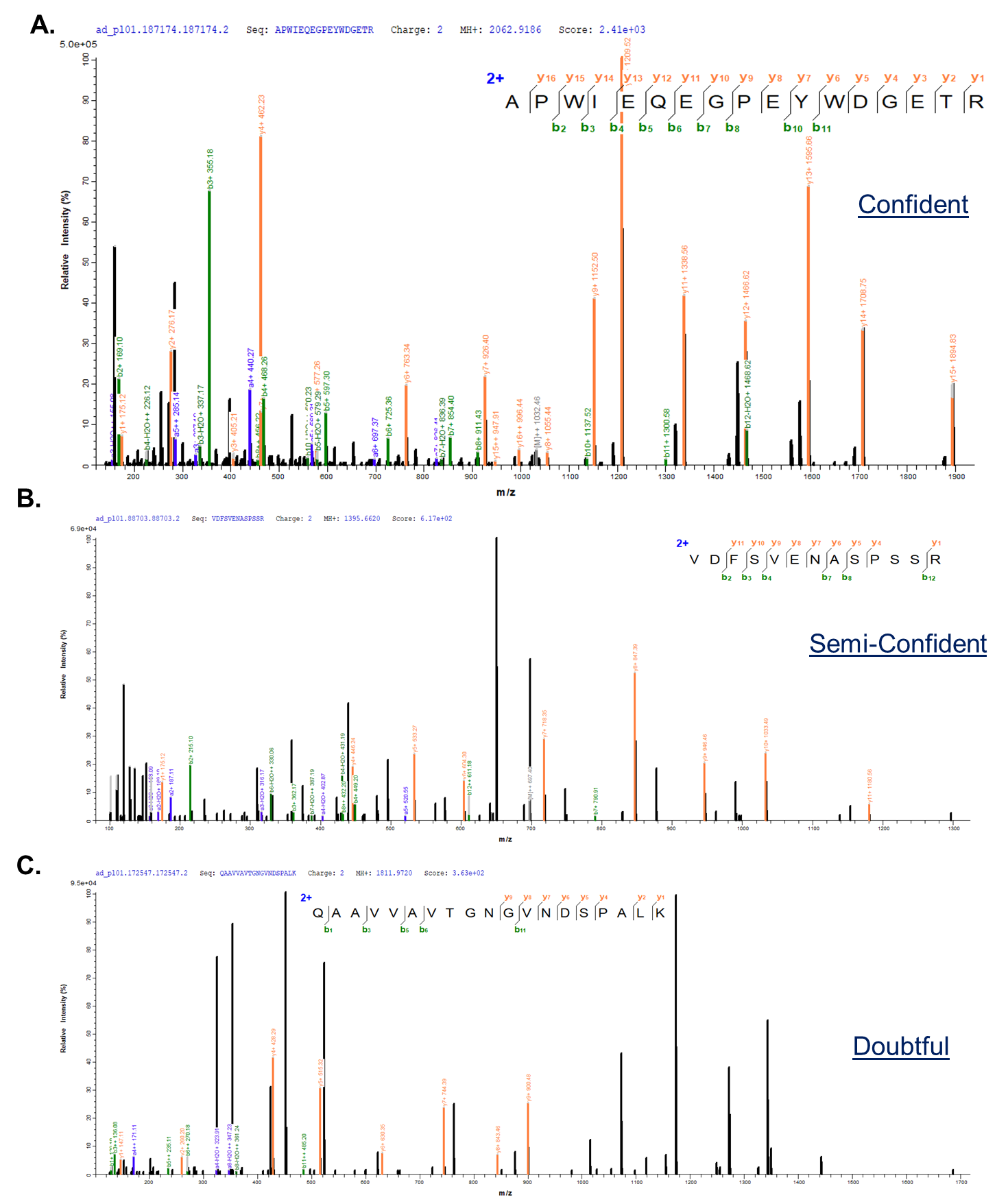

*Supplementary Figure S10: Examples for spectra classified with VAS in (A) confident, (B) semi-confident, (C) doubtful category.*

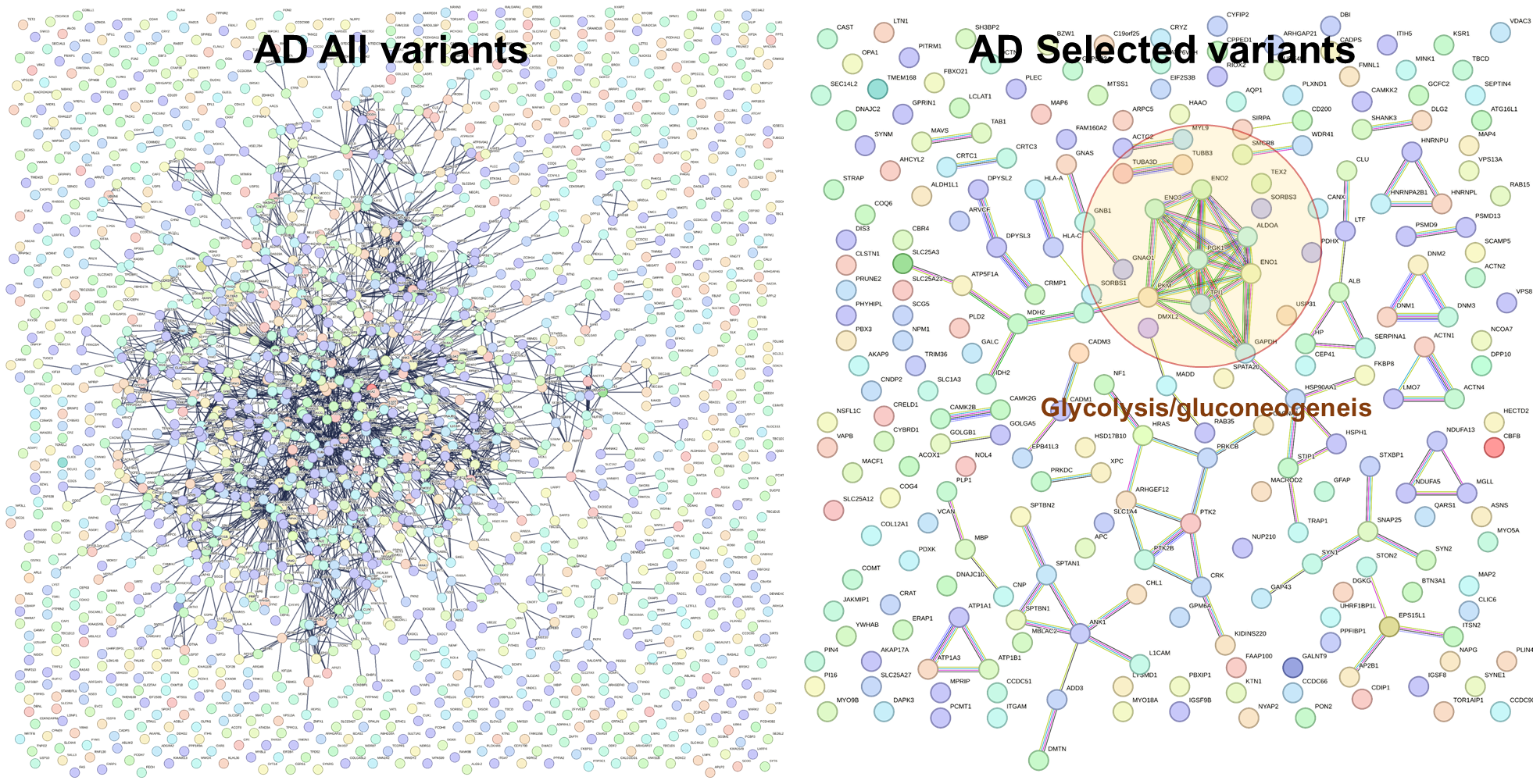

*Supplementary Figure S11: AD all variant genes (left) depict PPI network of all pathways in non-specific manner. When selected high quality variants are used, glycolytic/gluconeogenesis pathways are highlighted.*

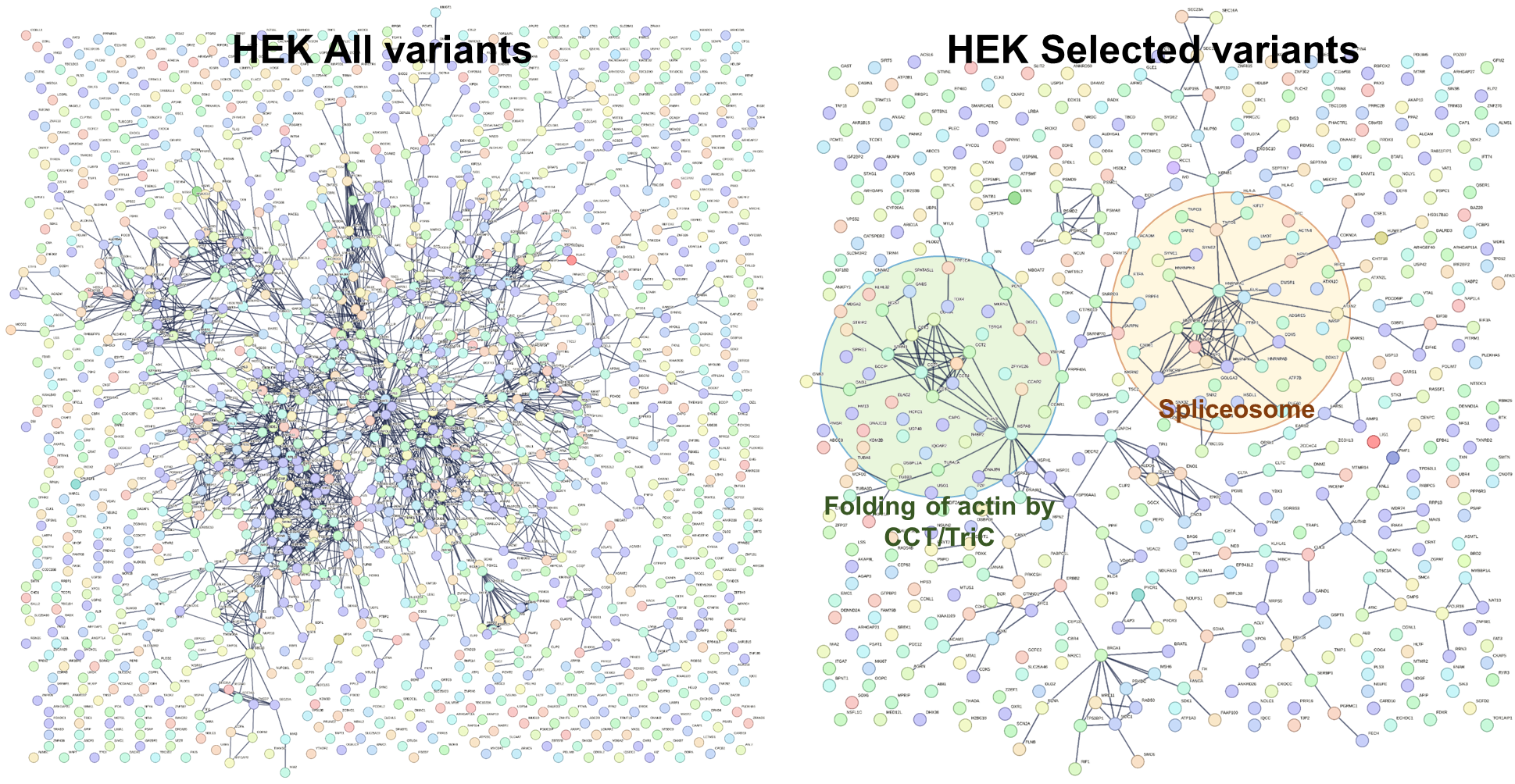

*Supplementary Figure S12: HEK all variant genes (left) depict PPI network of all pathways in non-specific manner. When selected high quality variants are used, spliceosomes and actin folding pathways by CCT/Tric are highlighted.*

**Supplementary Tables**

*Supplementary Table S1: MaSS-Simulator Parameters used for generation of simulated spectra*

| C-ions | Mass Offsets | 0 -1.0 +1.0 -18.015 -17.013 |
| --- | --- | --- |
|  | Ion Generation Probabilities | 100 70 70 70 70 |
|  | Ion Intensities | 1.75 1 1 0.25 0.25 |
| N-ions | Mass Offsets | 0 -1.0 +1.0 -18.015 -17.013 -28.01 -45.023 -46.025 |
|  | Ion Generation Probabilities | 100 70 70 70 70 70 70 70 |
|  | Ion Intensities | 1.5 1 1 0.75 0.75 0.25 0.25 0.25 |
| Immonium ions | Amino Acids | M D Q K E |
|  | Ion Generation Probabilities | 68 68 68 68 68 |
|  | Ion Intensities | 0.02 0.02 0.02 0.02 0.02 |
| Multiple charged ions | Charges | 2 |
|  | Ion Generation Probabilities | 35 |
| Noise settings | Noise Type | 2 |
|  | POS | 80 |
|  | Intensity Range | 0.02-1.0 |
| Other Settings | Precursor Mass Range | 500-20000 |
| Input File Formatting | Number of modifications allowed per peptide | 3 |
|  | % Probability that each peptide will contain a modification | 100 |
|  | Modification | C+ 57.02 |

*Supplementary Table S2: Composition of peptides used for simulation spectra generation (MS/MS) and sequences in the search database. The strategy behind the decisions of this design with identified, missed, and falsely identified PSMs are also provided.*

| Spectra | Database | Remarks | Identified Spectra | Missed Spectra | False Positives |
| --- | --- | --- | --- | --- | --- |
| Spectra with one variant: 5000 | 5000 Variant peptides  (1 variant) | True Positives | 4943 | 57 | 0 |
| Spectra with two variants: 5000 | 5000 Variant peptides  (2 variants) | True Positives | 4956 | 44 | 4 |
| Spectra without any variant (Wildtype): 40,000 | Sequence absent corresponding to these 40,000 peptides | False Positives  (To enhance the competition and make WT spectra hit on variant) | 0 | 38528 | 1472 |
| Unrelated variant spectra absent/not included | 5,00,000 Non-related variant peptides which are missing in simulated spectra | False Positives | - | - | - |

*Supplementary Table S3: VAS classification against manual validation classes for F1 fraction of AD dataset.*

| **All PSMs** | **Annotation** | **Manual** | | | **Total** |
| --- | --- | --- | --- | --- | --- |
|  |  | **Good** | **Average** | **Bad** |  |
| **VAS** | Confident | 175 | 85 | 10 | **270** |
|  | Semi-Confident | 16 | 66 | 17 | **99** |
|  | Doubtful | 30 | 212 | 186 | **428** |
|  | **Total** | **221** | **363** | **213** |  |

*Supplementary Table S4: PgxSAVy output on AD dataset comparing gFDR and cFDR.*

| AD dataset | | gFDR | | | cFDR | | |
| --- | --- | --- | --- | --- | --- | --- | --- |
| Fraction | Name used in this study | Confident | Semi-Confident | Doubtful | Confident | Semi-Confident | Doubtful |
| ad_pl01 | F1 | 232 | 98 | 398 | 270 | 99 | 428 |
| ad_pl02 | [F2](mailto:F@) | 234 | 93 | 479 | 251 | 101 | 568 |
| ad_pl03 | F3 | 287 | 163 | 720 | 297 | 186 | 850 |
| ad_pl04 | [F4](mailto:F@) | 101 | 94 | 632 | 98 | 91 | 669 |
| ad_pl05 | F5 | 135 | 76 | 476 | 150 | 88 | 547 |
| ad_pl06 | [F6](mailto:F@) | 225 | 75 | 423 | 238 | 88 | 490 |
| ad_pl07 | F7 | 233 | 51 | 346 | 233 | 51 | 346 |
| ad_pl08 | [F8](mailto:F@) | 221 | 99 | 517 | 230 | 101 | 639 |
| ad_pl09 | F9 | 36 | 102 | 565 | 54 | 102 | 676 |
| ad_pl10 | [F10](mailto:F@) | 218 | 51 | 355 | 263 | 80 | 493 |

*Supplementary Table S5: PgxSAVy output on HEK dataset comparing gFDR and cFDR.*

| **HEK Dataset** | | **gFDR** | | | **cFDR** | | |
| --- | --- | --- | --- | --- | --- | --- | --- |
| Datasets | Name used in this study | Confident | Semi-Confident | Doubtful | Confident | Semi-Confident | Doubtful |
| b1906_293T_proteinID_01A_QE3_122212 | F1 | 8 | 16 | 56 | 9 | 18 | 58 |
| b1922_293T_proteinID_02A_QE3_122212 | [F2](mailto:F@) | 54 | 17 | 71 | 57 | 17 | 74 |
| b1923_293T_proteinID_03A_QE3_122212 | F3 | 18 | 12 | 58 | 20 | 11 | 64 |
| b1924_293T_proteinID_04A_QE3_122212 | [F4](mailto:F@) | 47 | 11 | 76 | 53 | 19 | 96 |
| b1925_293T_proteinID_05A_QE3_122212 | F5 | 29 | 14 | 111 | 31 | 13 | 120 |
| b1926_293T_proteinID_06A_QE3_122212 | [F6](mailto:F@) | 44 | 11 | 59 | 45 | 14 | 67 |
| b1927_293T_proteinID_07A_QE3_122212 | F7 | 29 | 20 | 98 | 29 | 20 | 98 |
| b1928_293T_proteinID_08A_QE3_122212 | [F8](mailto:F@) | 34 | 11 | 89 | 35 | 11 | 92 |
| b1929_293T_proteinID_09A_QE3_122212 | F9 | 48 | 30 | 168 | 51 | 27 | 176 |
| b1930_293T_proteinID_10A_QE3_122212 | [F10](mailto:F@) | 37 | 13 | 67 | 43 | 20 | 72 |
| b1931_293T_proteinID_11A_QE3_122212 | F11 | 28 | 20 | 122 | 32 | 24 | 176 |
| b1932_293T_proteinID_12A_QE3_122212 | [F12](mailto:F@) | 28 | 7 | 49 | 42 | 13 | 95 |
| b1937_293T_proteinID_01B_QE3_122212 | F13 | 62 | 13 | 132 | 70 | 20 | 163 |
| b1938_293T_proteinID_02B_QE3_122212 | [F14](mailto:F@) | 26 | 9 | 68 | 27 | 12 | 128 |
| b1939_293T_proteinID_03B_QE3_122212 | F15 | 41 | 19 | 89 | 45 | 20 | 94 |
| b1940_293T_proteinID_04B_QE3_122212 | [F16](mailto:F@) | 39 | 3 | 72 | 40 | 16 | 144 |
| b1941_293T_proteinID_05B_QE3_122212 | F17 | 37 | 15 | 99 | 44 | 23 | 141 |
| b1942_293T_proteinID_06B_QE3_122212 | [F18](mailto:F@) | 45 | 13 | 131 | 47 | 16 | 146 |
| b1943_293T_proteinID_07B_QE3_122212 | F19 | 47 | 11 | 58 | 47 | 13 | 62 |
| b1944_293T_proteinID_08B_QE3_122212 | [F20](mailto:F@) | 40 | 23 | 153 | 41 | 22 | 156 |
| b1945_293T_proteinID_09B_QE3_122212 | F21 | 12 | 3 | 8 | 8 | 5 | 15 |
| b1946_293T_proteinID_10B_QE3_122212 | [F22](mailto:F@) | 58 | 25 | 129 | 62 | 47 | 191 |
| b1947_293T_proteinID_11B_QE3_122212 | F23 | 23 | 5 | 9 | 29 | 5 | 10 |
| b1948_293T_proteinID_12B_QE3_122212 | [F24](mailto:F@) | 28 | 5 | 16 | 38 | 18 | 121 |
